# Approximate Bayesian computation reveals the importance of repeated measurements for parameterising cell-based models of growing tissues

**DOI:** 10.1101/194712

**Authors:** Jochen Kursawe, Ruth E. Baker, Alexander G. Fletcher

**Affiliations:** Mathematical Institute, University of Oxford, Andrew Wiles Building, Radcliffe Observatory Quarter, Woodstock Road, Oxford, OX2 6GG, UK; Present address: Faculty of Biology, Medicine and Health, University of Manchester, Michael Smith Building, Oxford Road, Manchester, M13 9PL, UK; School of Mathematics and Statistics, University of Sheffield, Hicks Building, Hounsfield Road, Sheffield, S3 7RH, UK; Bateson Centre, University of Sheffield, Sheffield, S10 2TN, UK

**Keywords:** cell-based models, approximate Bayesian computation, parameter inference, vertex models, *Drosophila* wing imaginal disc

## Abstract

The growth and dynamics of epithelial tissues govern many morphogenetic processes in embryonic development. A recent quantitative transition in data acquisition, facilitated by advances in genetic and live-imaging techniques, is paving the way for new insights to these processes. Computational models can help us understand and interpret observations, and then make predictions for future experiments that can distinguish between hypothesised mechanisms. Increasingly, cell-based modelling approaches such as vertex models are being used to help understand the mechanics underlying epithelial morphogenesis. These models typically seek to reproduce qualitative phenomena, such as cell sorting or tissue buckling. However, it remains unclear to what extent quantitative data can be used to constrain these models so that they can then be used to make quantitative, experimentally testable predictions. To address this issue, we perform an *in silico* study to investigate whether vertex model parameters can be inferred from imaging data, and explore methods to quantify the uncertainty of such estimates. Our approach requires the use of summary statistics to estimate parameters. Here, we focus on summary statistics of cellular packing and of laser ablation experiments, as are commonly reported from imaging studies. We find that including data from repeated experiments is necessary to generate reliable parameter estimates that can facilitate quantitative model predictions.

## 1 Introduction

The growth and dynamics of epithelial tissues are central to numerous processes in embryonic development. Experimental studies are generating an increasing volume of quantitative and semi-quantitative data on these processes [1], providing the potential for significant increases in our understanding of epithelial morphogenesis. Computational models can be used to interpret such data, develop theories for the underlying biophysical mechanisms, and make predictions for future experiments to distinguish between competing theories. Increasingly, ‘cell-based’ modelling approaches are used that exploit the availability of data across subcellular, cellular and tissue scales [2]. Usually, these models are calibrated and tested according to their ability to reproduce qualitative phenomena and, as such, their predictive capability is limited. It remains largely unclear to what extent cell-based models can be used in a quantitative manner.

One important challenge that arises when applying cell-based models quantitatively is parameter inference from experimental data. Since cell-based models predict high-dimensional data, for example the shape or position of each modelled cell, it is necessary to select low-dimensional summary statistics to infer model parameters. However, the optimal choice of summary statistic or experimental design to infer parameters is non-intuitive. A common approach to parameter inference for computational models is to identify parameters of best fit. However, this approach is problematic since it does not quantify the uncertainty associated with such estimates. Here, we conduct parameter inference, compare summary statistics, and estimate uncertainty for a vertex model of the *Drosophila* wing imaginal disc, a classical model system for the study of tissue growth [3–6]. Vertex models are a widely used class of cell-based model in which epithelial cell sheets are approximated by tessellations of polygons representing the apical surfaces of cells, and vertices (where three or more cells meet) move in response to forces due to growth, interfacial tension and the hydrostatic pressure within each cell [5, 7, 8].

Many cell-based models include mechanical parameters that influence dynamic and steady-state behaviour. The optimal choice for such parameters for a given biological system is often not intuitive, since these parameters reflect mechanical properties of single cells or their pairwise interactions. To measure the mechanical properties of a single cell it may be necessary to remove it from the surrounding tissue, however this generally has a significant impact on its mechanical properties. Hence, parameterising cell-based models using tissue-scale data is an area of growing interest [9]. In the case of vertex models, efforts have been made to estimate mechanical parameters from tissue-scale measurements. Farhadifar et al. [5] achieve this by manual fitting of their model using a combination of summary statistics: the relative occurrence of cells with different numbers of sides; the average area of cells with different numbers of sides; the maximum displacement of vertices following laser ablation of individual cell edges; and the area and perimeter changes for cells whose edges were ablated. Other authors have used similar statistics to parameterise vertex models of the *Drosophila* wing imaginal disc [6, 10, 11] and the *Xenopus laevis* animal cap [12]. These authors arrive at different parameter estimates, which is not surprising given that the analysed tissues, summary statistics, and model implementations differ.

A variety of approaches have been taken to fit summary statistics obtained from vertex models to experimental data. These include ad hoc [10, 11] and least-squares [6, 12] approaches. Merzouki et al. [13] infer parameters by inducing tissue deformation; they compare stress-strain curves obtained from simulations with those obtained from experiments on monolayers of Madine-Darby Canine Kidney cells [14]. In these experiments, a free monolayer is suspended between rods and the stress curve is recorded as the distance between the rods was increased. A similar approach is taken by Xu et al. [15, 16], who highlight that the stress-strain curves obtained in this way are affected by the amount and orientation of cell divisions. Similarly, Wyatt et al. [17] show that cell divisions in this tissue are oriented so as to dissipate stress. Studies of cell mechanical properties in epithelia also include force-inference methods [18–20], where a heterogeneous generalisation of the vertex model is fitted to microscopy data to estimate the apical tension and pressure forces on each cell in the tissue. Such approaches are particularly suitable to estimate mechanical heterogeneity in a tissue without the need to physically manipulate the sample, though they typically assume the tissue to be in mechanical equilibrium, which is generally not the case during embryogenesis.

In summary, although some progress has been made in estimating the parameters of vertex models, none of the studies described above quantify the uncertainty of *in vivo* parameter estimates, nor do they address parameter identifiability. Hence, it remains unclear to what extent previously used summary statistics are sensible choices for parameter inference. Tissue manipulation approaches are not applicable *in vivo* since they require the removal of a tissue from its substrate [13–15], while force-inference methods do not directly estimate model parameters. There is thus a need to establish to what extent each parameter can be estimated, and the associated uncertainty quantified, in such models. For the first time we apply Bayesian inference to estimate vertex model parameters in a simulation of tissue growth. Bayesian inference enables uncertainty quantification by calculating the probability of the model parameters given observed data. Specifically, we apply Approximate Bayesian computation (ABC) [21, 22], a method that is commonly used when the likelihood of the model is not analytically tractable. To evaluate the inference method and compare summary statistics we apply the inference method to virtual data generated from the model, which allows us to compare inferred parameter values to the ground truth.

The remainder of this paper is structured as follows. In Section 2 we describe our *in silico* study and the inference method in detail. In Section 3 we infer model parameters using a range of summary statistics. We find that the uncertainty in parameter estimates generated using summary statistics of cell packing, or from tissue responses to laser ablations, depends heavily on the amount of data used to calculate summary statistics, and we investigate how much data is required for reliable parameter estimation. We identify the mean area of cells of each polygon class, in combination with the mean cell elongation, as a suitable summary statistic for vertex model parameter inference, and analyse how the quality of parameter estimates varies over parameter space. In Section 4 we discuss the implications of our findings.

## 2 Methods

### 2.1 Vertex model

For our *in silico* study, we consider a simplified vertex model of wing imaginal disc growth in the dipteran fly *Drosophila melanogaster*. The wing imaginal disc has been intensely studied as an experimental system for research on tissue growth and proliferation. During larval stages, this epithelial tissue undergoes a period of intense proliferation, increasing from around 40 to over 50,000 cells, before later giving rise to the adult wing [5, 23]. We now outline the technical details of our model; note that, motivated by the need to run a large number of simulations for parameter inference, we include several simplifying assumptions.

We represent the wing disc epithelium as a dynamic tessellation of polygons that approximate cell apical surfaces, with a vertex at each point where three cells meet. The position of each vertex, i, evolves in time according to the overdamped force equation

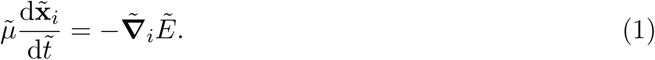

Here and throughout, we use tildes to denote dimensional quantities. In equation (1), 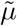 denotes the friction strength, 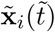 is the position of vertex *i* at time 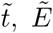 denotes the total energy associated with the tissue, and 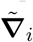 denotes the gradient operator with respect to 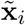. The number of vertices in the system may change over time due to cell division and removal (see below). For a homogeneous (unpatterned) epithelial tissue, the total energy 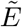 is given by [5]

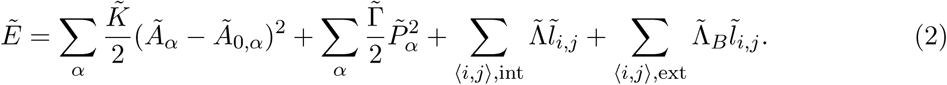

where the first two sums run over every cell *α* in the tissue, the third sum runs over every cell edge (pair of neighbouring vertices) ⟨*i*, *j*⟩ internal to the tissue and the third term runs over all cell edges at the boundary of the tissue, at time 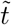. In equation (2), the variables *Ã*_*α*_ and 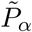 denote the area and perimeter of cell *α*, respectively, and the parameter *Ã*_0,*α*_ denotes a ‘target’ or preferred area for that cell. The four sums respectively represent the bulk elasticity of each cell, the presence of a contractile acto-myosin cable along the perimeter of each cell, and the combined effect of binding energy and contractile molecules at the interface between two cells or between cells and the tissue boundary. The parameters 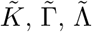 and 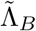 govern the strengths of each of these energy contributions.

Before solving the model numerically, we non-dimensionalise it to reduce the number of free parameters [5]. Rescaling space by a characteristic cell length scale, 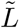, and time by a characteristic timescale, 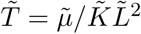, we obtain

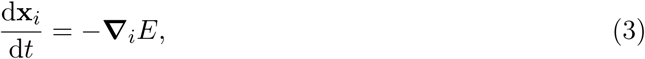

where

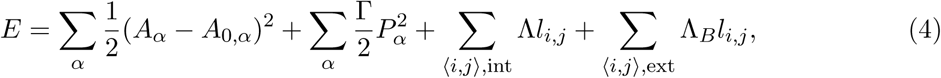

and

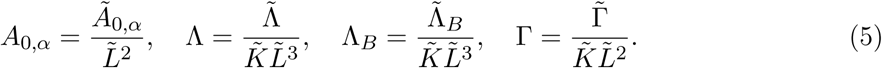

We solve equations (3) and (4) numerically using the forward Euler scheme

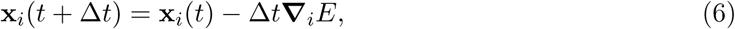

where Δ*t* = 0.01 is chosen to be sufficiently small to guarantee convergence [24].

Initially, the tissue is represented by four hexagonal cells of unit area (Figure 1A). The initial target area of these cells 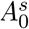 is the same as their area, 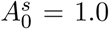. The boundary of the tissue is allowed to move freely throughout the simulation. During our simulations we allow cell neighbour exchange through T1 transitions (Figure 1B) and cell removal through T2 transtions (Figure 1C). T1 transitions are executed whenever the length of a given edge decreases below the threshold *l*_T1_ = 0.01. The length of the new edge, *l*_new_ = *ρl*_T1_ (*ρ* = 1.5), is chosen to be slightly longer than this threshold to avoid an immediate reversion of the transition.

**Figure 1:**
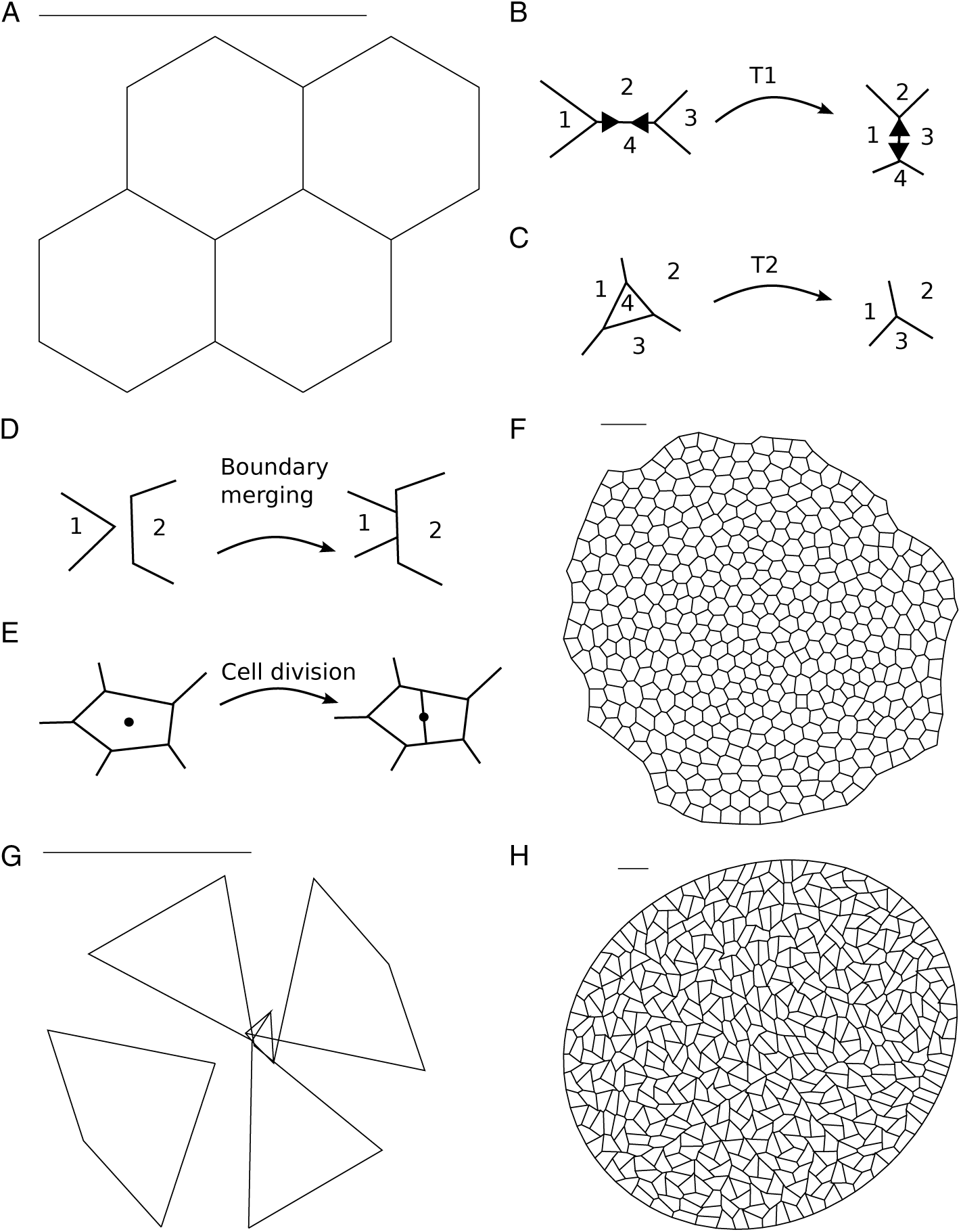
Simulation setup and boundary conditions. (A) The initial condition comprises four hexagonal cells. (B-E) Vertex configurations can change through T1 swaps (B), T2 swaps (C), boundary merging (D) and cell division (E). (F) The tissue grows for *n*_*d*_ = 7 rounds of divisions until it comprises approximately 500 cells. (G) If the line tension parameter, Λ, is negative in the bulk of the tissue as well as on the tissue boundary the tissue may assume unphysical configurations. (H) We prevent such artefacts by setting the line tension along the tissue boundary to zero when Λ is negative, leading to well-defined tissue geometries. Panels (A,F-H) are rescaled to fit the view; a scale bar is added for comparison.

During a T2 transition, a small triangular cell or void is removed from the tissue and replaced by a new vertex. In our implementation any triangular cell is removed if its area drops below the threshold *A*_T2_ = 0.001. The energy function (4), in conjunction with T2 transitions, can be understood as a model for cell removal: cells are extruded from the sheet by a T2 transition if the energy function (4) leads to a sufficiently small cell. Note that in equation (4) the bulk elasticity or area contribution of a cell *α* is finite even when the area *A*_*α*_ is zero, allowing individual cells to become arbitrarily small if this is energetically favourable. As cells decrease in area they typically also reduce their number of sides. Hence, it is sufficient to remove only small triangular cells instead of cells with four or more sides [5, 6, 23].

We further model the merging of overlapping tissue boundaries (Figure 1D). Whenever two boundary cells overlap, a new edge of length *l*_new_ is created that is shared by the overlapping cells. In cases where the cells overlap by multiple vertices, or if the same cells overlap again after a previous merging of edges, the implementation ensures that two adjacent polygons never share more than one edge by removing obsolete vertices. The merging of boundary edges is discussed in further detail in [7].

From the initial condition shown in Figure 1A, we simulate *n*_*d*_ = 7 rounds of cell division in the tissue. Following previous work [10, 24], we assume that each cell proceeds through two cell cycle phases: interphase and growing. Letting *t*_cycle_ denote the average total cell cycle duration, we assume that the time spent in interphase is drawn independently from an exponential distribution with mean 2*t*_cycle_/3. We introduce stochasticity in this phase to prevent unrealistically synchronous daughter cell divisions. After this age, the cell experiences a growing phase of fixed duration *t*_cycle_/3. During this time the cell’s target area, *A*_0,*α*_, grows linearly to twice its original value. Upon completion of the growth phase, the cell divides.

During cell division, a new edge is created that separates the newly created daughter cells (Figure 1E). The new edge is drawn along the short axis of the polygon representing the parent cell [7, 25]. Daughter cells are initialised with target area 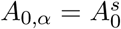, which corresponds to half the target area of the mother cell upon division.

Note that the precise number of cells at the end of each simulation differs due to variations in the number of cell removals through T2 transitions. Each cell of the last generation remains in interphase until the simulation stops. We select the total simulation time to be *t*_end_ = 700, which is sufficiently long for the tissue to settle into a static configuration after the last rounds of division. This is illustrated in Supplementary Figure S1.

We implement the model within Chaste, an open source C++ library that provides a systematic framework for the simulation of vertex models [7, 24, 26]. Our code is available in the supplementary material as a zip archive. Each time step starts by updating the cell target areas. Cell divisions, removals, neighbour exchanges, and boundary merging are then performed before updating the vertex positions and incrementing the simulation time by Δ*t*. The algorithm halts when the time reaches *t*_end_ = 700.

The parameter Λ_*B*_ is used to calculate the line tension of cell interfaces with the tissue boundary, i.e. the third term in equation (4). If the line tension parameter Λ is negative, we set the line tension of edges along the boundary to zero, Λ_*B*_ = 0. Otherwise, we set Λ_*B*_ = Λ. This boundary condition helps prevent boundary artefacts that lead to unphysical tissue shapes: in Figure 1G, the simulation parameters are Λ = −0.85 and Γ = 0.1, and the line tension for cell edges along the tissue boundary is the same as throughout the tissue, i.e. Λ_*B*_ = Λ = −0.85. In simulations with this parameter set, cells disconnect from each other or self-intersect, leading to failure of the simulation algorithm. In contrast, for simulations using Λ = −0.85, Γ = 0.1, Λ_*B*_ = 0, the tissue grows normally and leads to a physically meaningful tissue configuration (Figure 1H). If Λ is negative, the energy contribution of individual edges decreases as the edge length increases. Since the motion of boundary vertices is unopposed by neighbouring cells, cells at the boundary may thus grow to arbitrarily large sizes or self-intersect if the associated gain in energy from the edges is sufficiently large. Such boundary effects are prevented if Λ_*B*_ = 0. An adjustment of the boundary tension is not necessary if Λ is positive, since in this case it is energetically favourable for edges to shorten.

The non-dimensionalised parameter values used in our simulations are listed in Table 1. These parameter values are chosen so that the final tissue is sufficiently large to obtain tissue-level summary statistics while minimising the amount of time that is required to run a single simulation. Ensuring that individual simulations are quick to run is a prerequisite for conducting ABC, since a large number of samples are required from the prior distribution. We discuss simulation times further later in this section.

**Table 1:**
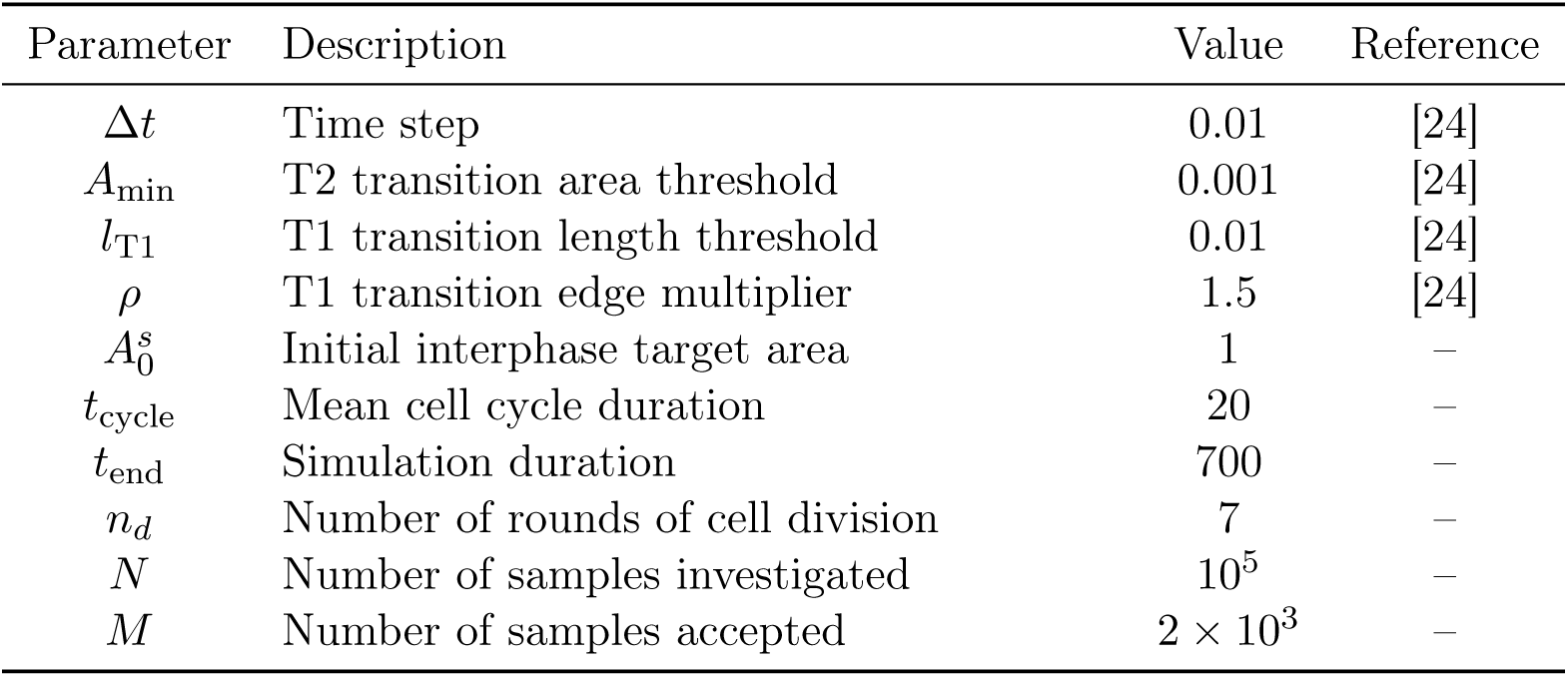
Description of non-dimensionalised parameter values.

Kursawe et al. [24] found that a key descriptor of cell packing, the average area of cells of each cell neighbour (or polygon) number, was unaffected by changes in the cell cycle duration over multiple orders of magnitude. This suggests that characteristics of cell packing in the final tissue configuration are not affected by an acceleration of tissue growth and that static tissue configurations in the vertex model are largely governed by the energy equation (4) and are less strongly affected by the dynamic processes leading up to the final configuration.

To estimate to what extent our implementation of the model may influence tissue properties, we compare outcomes between our simulation procedure and the one described by [5] for three points in parameter space in Figure 2. We find that cell packings generated using the model described here closely resemble those simulated previously by [5]. Further, the average area per polygon class (a sample summary statistic of cell packing) is similar to those previously reported for all three considered parameter combinations. In Figure 2D, the mean area per polygon class deviates between our simulations and those reported by Farhadifar et al. [5] for seven-and eight-sided cells. This difference may originate from differences in boundary conditions between our simulations and those by [5], or from differences in the description of cell growth and division. Overall, however, we conclude that the final vertex configuration in our model represents typical cell packing geometries of the vertex model for different points in (Γ, Λ) parameter space.

**Figure 2:**
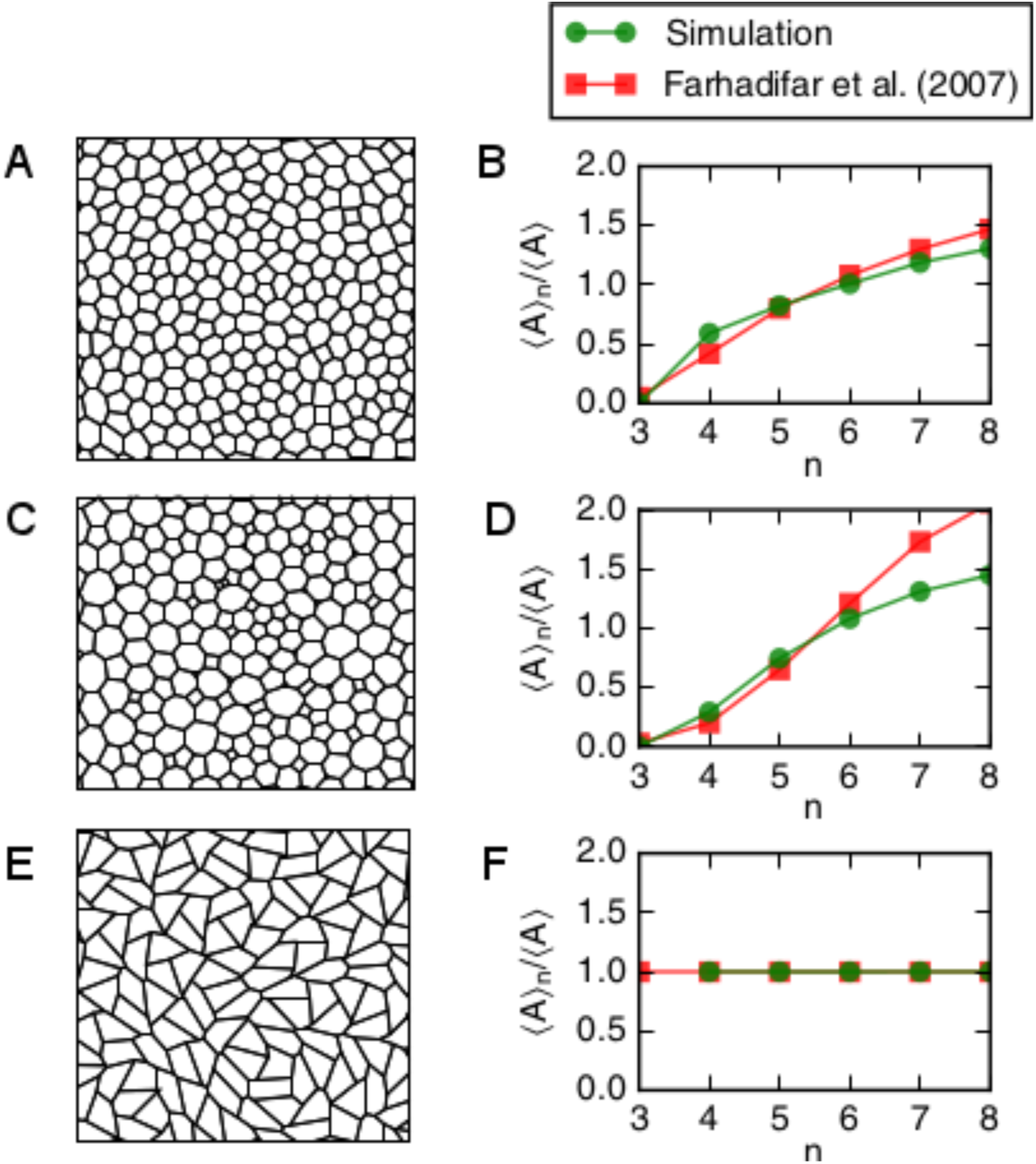
Cell packings depend on the vertex model parameters Λ and Γ. (A,C,E) Subsections of the tissue at the end of the simulation for different parameter values. Cell shapes differ between the parameter values. (C,F,I) Cell shapes are quantified by the mean area of cells of each polygon class, and compared between our simulation outcomes and those by [5]. Parameter combinations are: (Λ, Γ) = (0.12, 0.04) (A,B); (Λ, Γ) = (0.0, 0.1) (C,D); and (Λ, Γ) = (-0.85, 0.1) (E,F).

### 2.2 Parameter inference

In Bayesian statistics, one considers the joint probability distribution *p*(**Θ**, ***D***) of a parameter vector **Θ** and observed data vector ***D*** in order to calculate the posterior distribution *p*(**Θ**|***D***), the probability distribution of the parameters given the data. The calculation of the posterior is achieved by applying Bayes’ rule

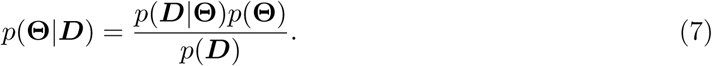

Here, *p*(***D***|**Θ**) is the probability of ***D*** given **Θ**, and is usually referred to as the likelihood, and *p*(***D***), the probability of observing the data, is the marginal likelihood. The likelihood of the parameter *p*(**Θ**) is referred to as the prior.

Approximate Bayesian computation (ABC) [21, 22] is increasingly used to estimate parameters of complex models for which the likelihood is not analytically or computationally tractable. The most simple form of ABC, rejection-based ABC [21], approximates the posterior through random sampling of parameters from the prior and evaluating the model for each sample. Summary statistics are used in order to compare the model with data. This is necessary since the modelled data is high-dimensional: in case of vertex models it comprises the position of each modelled vertex and the cell adjacencies. Samples for which a chosen summary statistic is sufficiently close to the observed data are considered samples of the posterior, otherwise they are rejected. Various well-established adaptations exist, including Markov Chain Monte Carlo approaches [27] and sequential Monte Carlo techniques [28]. Here, we apply rejection-based ABC with regression adjustment, which leads to better estimates of the posterior than rejection sampling alone [21]. In addition, unlike more sophisticated methods, rejection-based ABC allows us to use a fixed set of sample simulations to evaluate the suitability of a broad range of summary statistics of the data at different points in parameter space.

#### 2.2.1 Calculating a suitable prior distribution

The parameter space of the vertex model with energy (4) has been explored previously [5, 29] and can be subdivided into three regions, labelled A, B, and C in Figure 3. In region A, characterized by zero shear-stress, stable minima of (4) do not exist since infinitely many configurations are possible at the energy minimum. For parameter values in this region, our simulations tend to halt prematurely due to self-intersecting or overlapping cells, in agreement with previous results [5, 12, 29]. In region B, well-defined energy minima exist, and cell packings vary as the parameters are changed within this region (Figure 2). In region C, cell areas vanish at the energy minimum, with simulated tissues progressively shrinking and cells being removed through T2 transitions. Thus, since regions A and C yield unphysical model behaviours, we define our prior to be a uniform distribution on region *B*, restricted to the domain Λ ≥ −1.5, 0 ≤ Γ ≤ 0.2. By choosing Γ ≤ 0.2 we follow previous work in ensuring that a sufficiently large region of parameter space can be explored in a reasonable amount of time [5, 12, 29].

**Figure 3:**
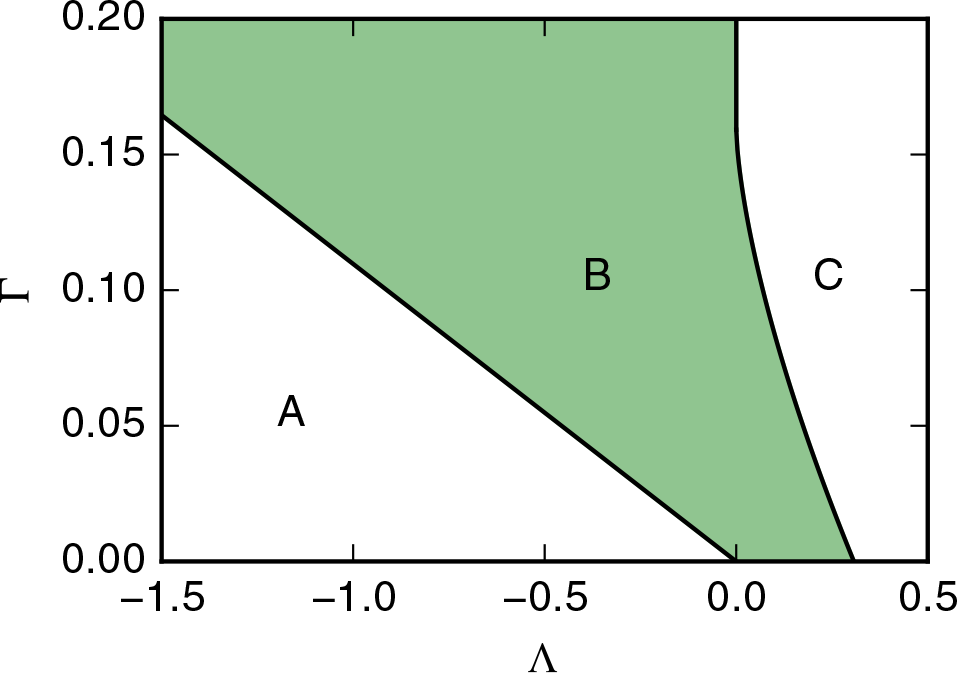
Definition of the prior. The parameter space of our vertex model can be divided into three regions [5, 29]. Region A marks configurations of zero shear stress, where infinitely many vertex configurations are possible at the energy minimum of equation (4). In region B, vertex configurations can be stable, corresponding to well-defined energy minima. In region C, the energy is minimal when all cells have zero area and zero perimeter. Throughout this work, the prior is chosen to be a uniform distribution over region B.

The borders of region B can be estimated by calculating the energy minima of equation (4) for tissues containing single, regular *n*-gons [29]. Different polygon classes give rise to different boundaries for the transition between stable configurations in region B and shear-free configurations in region A. Among all polygon classes, this boundary occupies the ‘left-most’ position for triangular cells, which allow the lowest values of Λ. For triangular cells, the boundary between regions A and B is given by the line [29]

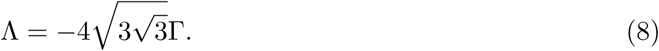

The boundary between regions B and C assumes positions at increasing values of Λ as the polygon number increases. As *n* → ∞, this position converges to [29]

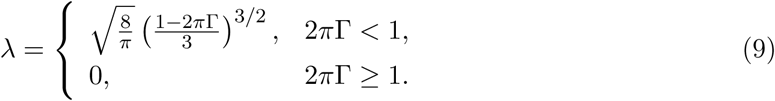

Using equations (8)–(9) we thus define the prior to exclude regions in parameter space where all simulations are unphysical^1^. However, unphysical solutions may also occur in region B of Figure 3 for parameters that are close to the boundaries (8) and (9).

#### 2.2.2 Sample generation

Within region B we generated *N* = 10^5^ uniformly distributed parameter samples and used these to run the *in silico* wing disc simulations using a supercomputer. Using the parameter values listed in Table 1, a single simulation requires approximately five minutes to run on a single machine with an Intel(R) Core(TM) i5-5200U CPU with a 2.20GHz processor and 8GB memory. Out of the 10^5^ parameter samples generated from the prior, 10,762 simulations did not run to completion due to unphysical events, for example the generation of self-intersecting or overlapping cells.

We further generated 10^5^ uniformly distributed parameter samples from the prior, and for each of these ran ten *in silico* wing disc simulations using a supercomputer. Of these repetition samples, 21,583 samples were incomplete due to unphysical events in one of the repetition simulations. Generating all samples required approximately 830,000 hours of calculation time.

#### 2.2.3 Implementation of ABC regression

In Section 3 we investigate the utility of a broad range of summary statistics in inferring vertex model parameters using *in silico* data generated using the procedure described in Section 2.1. Specifically, we evaluate the model at a reference parameter vector 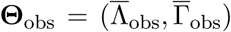. The samples used for parameter inference using ABC have parameter vectors 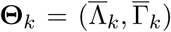, *k* = 1,…, *N*. A given vector of summary statistics computed with **Θ**_obs_ takes the vector form *s*_obs_ = (*s*_1_, …, *s*_*S*_)^T^ with a total of *S* components, and the summary statistic at sample **Θ**_*k*_ assume the values *s*_*k*_ = (*s*_*k*1_, …, *s*_*kS*_)^T^. Throughout we use summary statistics where the individual vector entries *s*_1_, …, *s*_*S*_ have been rescaled by their respective standard deviations across all samples. This is a common procedure in ABC, since it reduces the impact of vector components with high variance on the parameter estimate [21, 22].

Parameter inference is then conducted by first accepting the *M* = 2, 000 closest samples based on an Euclidean distance measure to the reference statistic, *d*_*i*_ = ||*s*_*i*_ - *s*_obs_||_2_. This accepted proportion is sufficiently small for all accepted samples to be close to the observed summary statistics, avoiding the need to explicitly select an appropriate acceptance distance. The acceptance threshold, *δ*, can be identified as the maximal distance of all accepted samples

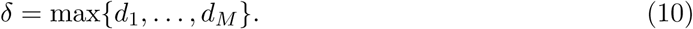

We next adjust the accepted parameter vectors **Θ**_*i*_ = (Λ_*i*_, Γ_*i*_) using local-linear regression, as introduced by [21]. Specifically, we minimise the penalty function

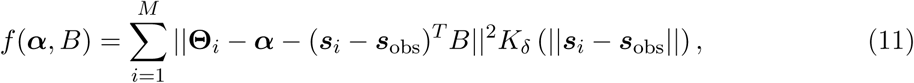

providing the regression parameters 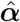 and 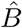 for a local-linear regression of the parameters dependent on the summary statistics,

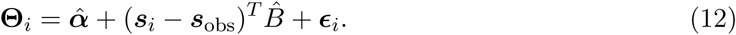

Here, *K*_*δ*_ denotes the Epanechnikov kernel function [30]

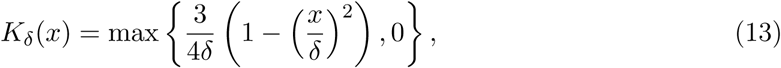

where *δ* is defined in equation (10) and the *ϵ*_*i*_ are uncorrelated random variables with zero mean and a common variance. Note that no further assumptions are made as to the distributions of *ϵ*_*i*_. The penalty function (11) is minimised by

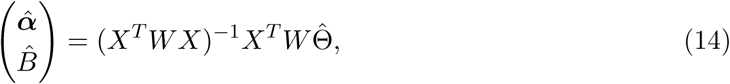

where the matrix *W* is diagonal with non-zero entries *K*_*δ*_(||*s*_*i*_ - *s*_obs_||), the matrix *X* has the form

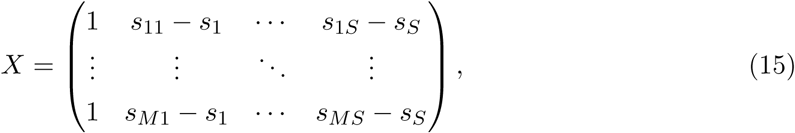

and the matrix 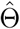 is given by (**Θ**_1_, …, **Θ**_*M*_)^*T*^.

After computing 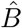 from equation (14) we perform regression adjustment on all samples,

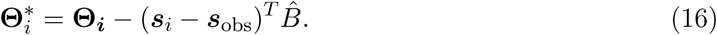

The process of regression adjustment aims to reduce the effects of the discrepancy between *s*_*i*_ and *s*_obs_. The vector 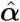 estimates the mean of the posterior distribution:

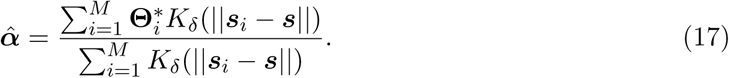

To estimate the posterior distribution, we use kernel density estimation with the Epanechnikov kernel. Specifically, each parameter vector 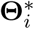 is assigned the weight *w*_*i*_ = *K*_*δ*_(||*s*_*i*_ - *s*_obs_||), and the posterior density at parameter 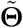 is calculated as

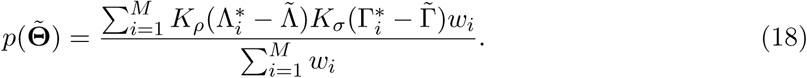

Note that when using kernel-density estimation, the estimated density at a given parameter point depends on the thresholds *ρ* and *σ* (whose values, we emphasise, may differ from that of *δ* in equation (10)). Throughout this work, we use least-squares cross-validation to select *ρ* and *σ* automatically [31].

#### 2.2.4 Definition of summary statistics

Vertex positions in our model are measured in units of the characteristic length scale of the tissue, which we set equal to the square root of the target area of cells in the tissue, 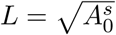. Since a cell’s target area may differ from its actual area, the target area is not experimentally accessible. Throughout this work we thus consider dimensionless summary statistics. Specifically, we divide all area-based summary statistics by the average area of cells in the tissue [12], and all length-based summary statistics by the square root of the average area of cells. A full list of all summary statistics is provided in Table 2. All summary statistics of cell shapes are averaged over all cells in the tissue that do not share an edge with the tissue boundary, since cell shapes along the tissue boundary differ from those in the bulk of the tissue (cf. Figure 1F,H). We define cell elongation as the square root of the ratio of the largest to the smallest eigenvalues of the moment of inertia of the polygon representing the cell. This definition provides a robust measure of elongation for arbitrary cell shapes [7].

**Table 2:** Description of summary statistics used in this study.

| Abbreviation | Summary statistic | Description |
| --- | --- | --- |
| Polygon numbers | $(p_4, \dots, p_8)$ | Cell neighbour number distribution |
| Area ratios | $(\langle A \rangle_4 / \langle A \rangle, \dots, \langle A \rangle_8 / \langle A \rangle)$ | Mean area for cells of each polygon class |
| Cell perimeter | $\langle P \rangle / \sqrt{\langle A \rangle}$ | Average cell perimeter |
| Edge length | $\langle l_{i,j} \rangle / \sqrt{\langle A \rangle}$ | Average edge length |
| Cell elongation | $\langle \lambda \rangle$ | Average cell elongation (see text for details) |
| Area deviation | $\sigma(A) / \langle A \rangle$ | Standard deviation of the cell areas |
| Area correlation | $\text{corr}(A_i, A_j)$ | Correlation between areas of adjacent cells $i$ and $j$ |
| Polygon number correlation | $\text{corr}(N_i, N_j)$ | Correlation between neighbour numbers of adjacent cells $i$ and $j$ |
| Neighbour areas | $(\langle A_{\text{neigh}} \rangle_4 / \langle A \rangle, \dots, \langle A_{\text{neigh}} \rangle_8 / \langle A \rangle)$ | Mean neighbour area for cells of each polygon class |
| Neighbour numbers | $(\langle N_{\text{neigh}} \rangle_4, \dots, \langle N_{\text{neigh}} \rangle_8)$ | Mean neighbour number for cells of each polygon class |
| Laser recoil | $\langle d_{\text{recoil}} \rangle / \sqrt{\langle A \rangle}$ | Recoil distance of vertices adjacent to an ablated edge |
| Area asymmetry | $\langle \Delta A \rangle / (d_{\text{recoil}} \sqrt{\langle A \rangle})$ | Relative change in area of the adjacent cells over the vertex recoil |
| Perimeter asymmetry | $(\langle \Delta P \rangle - 2d_{\text{recoil}}) / d_{\text{recoil}}$ | Relative change in perimeter of the adjacent cells over the vertex recoil |

Note that $p_n$ denotes the proportion of cells of polygon class $n$ ; $A_{\text{neigh}}$ and $N_{\text{neigh}}$ denote the average area of a cell's neighbours and the average polygon number of a cell's neighbours, respectively; and $\langle \cdot \rangle_n$ and $\langle \cdot \rangle$ denote averages over all cells of polygon class $n$ , and all cells, respectively, both excluding cells on the tissue boundary. For 'Polygon numbers', 'Area ratios', 'Neighbour areas' and 'Neighbour numbers' only cells of polygon classes four to eight are considered, since cells with different neighbour numbers are rarely observed in simulations.

Throughout this work we calculate correlations between two random variables *A* and *B* by

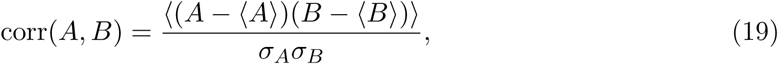

where *σ*_*X*_ denotes the standard deviation of a random variable *X*. In the case of ‘Area correlation’ and ‘Polygon number correlation’, the random variables are the areas and polygon numbers of adjacent cells, respectively. In these cases, the average in equation (19) is taken over all pairs of adjacent cells.

## 3 Results

In the following we address a key problem that arises when using cell-based models to interrogate quantitative experimental data: the practical identifiability of model parameters, and the choice of summary statistics for parameter inference and uncertainty estimation. We apply ABC for the first time to estimate mechanical parameters of a vertex model of a proliferative epithelium. We compare parameter estimates obtained using a range of summary statistics of our model and estimate the associated uncertainties.

When investigating the efficacy of different summary statistics in inferring the parameters of our model we use *in silico* data generated at the reference parameter set (Λ_obs_, Γ_obs_) = (0.12, 0.04). These are commonly used parameter values for vertex model simulations of the growing *Drosophila* wing imaginal disc [5, 23]. We begin by analysing the quality of the posterior distributions of Λ and Γ generated using different summary statistics of cell packing and tissue responses to perturbations. Specifically, we focus on first-order summary statistics that characterize the shapes of individual cells, second-order summary statistics of cell packing that quantify the relationships between shapes of adjacent cells, and summary statistics characterising the tissue response to perturbations through laser ablation experiments.

### 3.1 Parameter estimates from single experiments have high uncertainty

In Figure 4 we investigate the efficacy of various summary statistics obtained from single experiments in estimating vertex model parameters; posterior distributions obtained using first-order summary statistics, second order summary statistics, and summary statistics from laser ablation experiments are plotted. All of the investigated summary statistics are associated with high parameter uncertainty. In the following we provide background to each of these summary statistics and discuss the parameter estimates.

**Figure 4:**
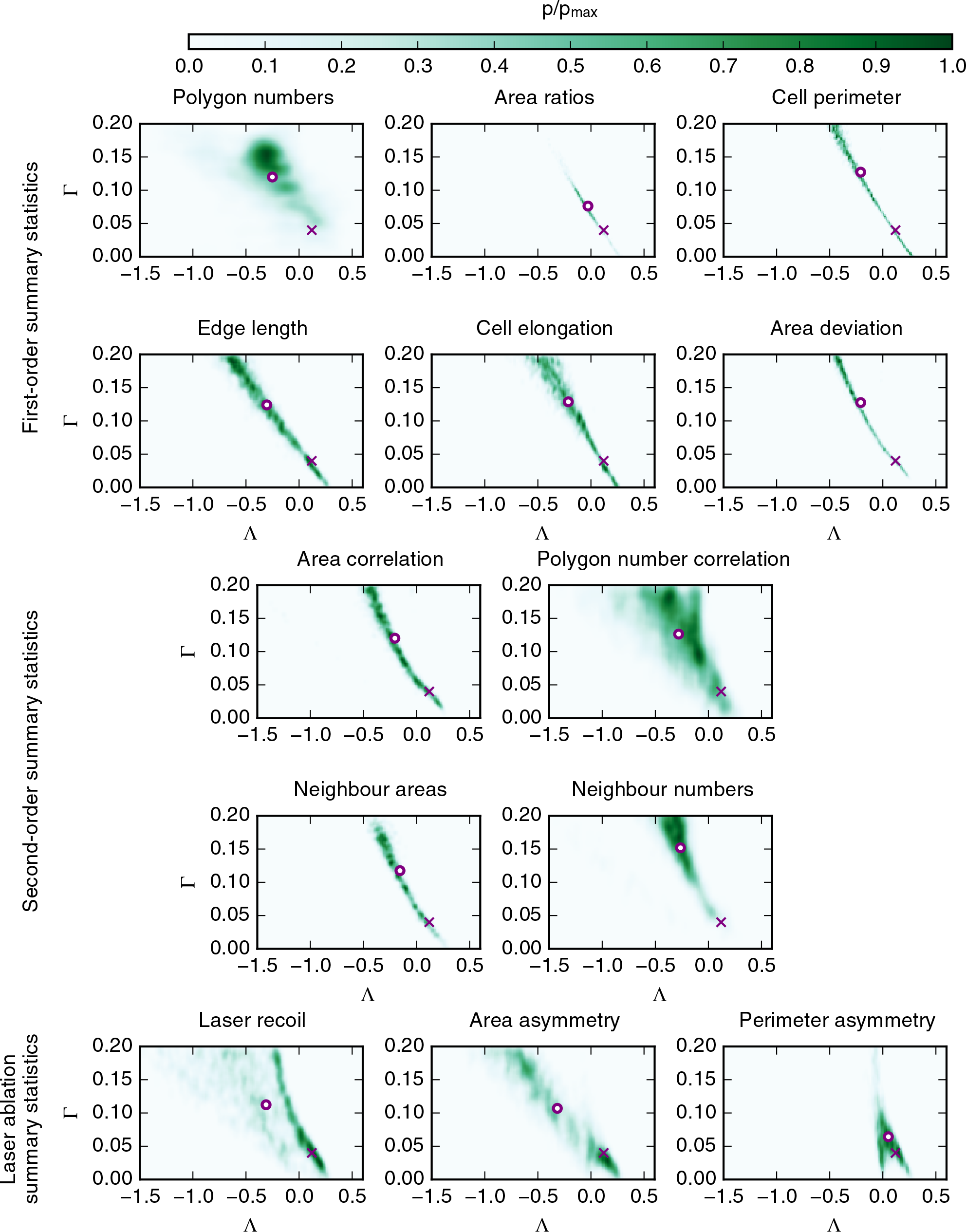
Posterior distributions of (Λ, Γ) obtained from different summary statistics of single experiments. Posterior distributions obtained using first-order summary statistics, second-order summary statistics, and laser ablation experiments are shown in green. Parameter values are listed in Table 1 and the summary statistics are defined in Table 2. Crosses mark the reference parameter set used to generate the observed data. Circles mark the means of the posterior distributions. Posterior distributions were calculated using summary statistics from single experiments.

First-order summary statistics of cell packing are often used to compare the output of vertex models with experimental data. The most common examples of these are the distribution of polygon (cell neighbour) numbers and the average area of cells for each polygon class. Here, we employ both of these summary statistics for parameter inference, in addition to the average cell perimeter, the average edge length (i.e. the average length of cell-cell interfaces), the average cell elongation, and the standard deviation of cell areas within the tissue.

The approximate posterior distributions generated by ABC using these first-order summary statistics are shown in Figure 4. The means of these posterior distributions provide an estimate of the inferred parameters and are calculated using equation (17). We find that none of the summary statistics considered lead to posterior distributions that accurately estimate parameters of the *in silico* data. The most accurate parameter estimate is achieved using the summary statistic ‘Area ratios’, with a parameter estimate of (Λ, Γ) = (-0.28, 0.076). The reference parameter set used to generate the *in silico* data is (Λ_obs_, Γ_obs_) = (0.12, 0.04). For the ‘Area ratios’ summary statistic, the posterior distribution is also the most narrow and concentrated, i.e. it is the posterior distribution with the lowest degree of uncertainty. The marginal standard deviations are 0.14 and 0.035 in Λ and Γ, respectively. All of the posterior distributions corresponding to first-order summary statistics in Figure 4 are spread across parameter space instead of concentrating in an area close to the true parameter set. Thus, the parameter estimates are all associated with a high degree of uncertainty, and therefore without knowing the reference parameter set it would be difficult for us to evaluate the quality of parameter estimates, including the cases of the commonly used polygon distribution and the cell area ratios. This could mean that the parameters are not practically identifiable using these summary statistics [32], or that insufficient data are used for the inference.

We next investigate the efficacy of second-order summary statistics in estimating vertex model parameters (Figure 4, middle). We focus on four quantities: the ‘Area correlation’, i.e. the correlation coefficient between areas of adjacent cells; the ‘Polygon number correlation’, i.e. the correlation coefficient between neighbour numbers of adjacent cells; the ‘Neighbour areas’, i.e. the mean area of neighbours for cells of each polygon class; and the ‘Neighbour numbers’, i.e. the mean polygon number of neighbours for cells of each polygon class.

In general, the second-order statistics considered in Figure 4 suffer the same drawbacks as the first-order summary statistics. In particular, they all yield posterior distributions covering wide regions in parameter space and the posterior means lie far away from the reference parameter set.

The shapes of the posterior distributions generated using a range of first-and second-order statistics in Figure 4 have striking similarities. For many of the summary statistics, such as ‘Area correlation’ and ‘Cell elongation’, the posterior distributions are spread along a narrow, elongated region in parameter space. This indicates that cell shapes and neighbour relationships generated by the vertex model within such regions are similar and indistinguishable by these summary statistics. The reason for this might be that Λ and Γ have distinct but related roles in the vertex model. The parameter Λ regulates the strength of an energy contribution that is linear in the edge length, and Γ regulates the strength of an energy contribution quadratic in the perimeter. Thus, both parameters affect the overall contraction force along cell-cell interfaces, which might explain why stable configurations in the vertex model appear similar in regions of decreasing Λ and increasing Γ. This relationship can be illustrated by considering the stability conditions for tissues containing single cells derived by Staple et al. [29], who considered tissues containing single polygons of target area 1, area *a*, and perimeter *p*. Such polygons are energetically stable if

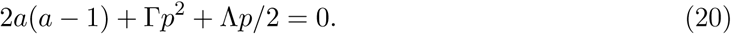

If a polygon with area *a* and perimeter *p* is stable for fixed values of Λ and Γ, then there are infinitely many other parameter combinations for which equation (20) is fulfilled for the same values of *a* and *p*. These parameter combinations all lie along the straight line in (Λ, Γ) parameter space,

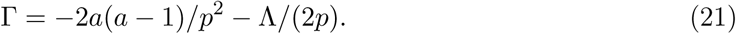

The shapes of the posterior distributions from first-and second-order statistics in Figure 4 indicate this rule may generalise to energy minima of tissues containing multiple cells.

Using first-and second-order summary statistics of cell shapes in Figure 4 we had limited success in inferring Λ and Γ reliably, revealing a possible interdependence of these parameters. It is intuitive that such static image-based methods have limitations, since they use cell shapes as a means to infer mechanical properties, and one may assume that invasive methods that record the dynamics of tissue response to a manipulation may produce more reliable results for the inference of mechanical parameters.

To test whether localised mechanical perturbation can be helpful in inferring parameters we then simulated laser ablation experiments. These are widely used to investigate mechanical properties of epithelial cells and comprise ablating the bond between two adjacent cells, usually leading the shared vertices of that edge or bond to recoil. Summary statistics of this recoil process can give insights into the mechanical state of the tissue or the adjacent cells [5, 33]. For each sample simulation we select one cell-cell interface in at the centre of the final tissue configuration and simulate a laser ablation of this interface. To simulate laser ablation, we remove the line tension contribution in equation (4) from the ablated edge and the perimeter contribution from both of its adjacent cells, and further set the threshold for vertex rearrangement 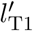 throughout the tissue to zero. The latter prevents vertex rearrangements in the cells adjacent to the ablation, which we consider physically unrealistic since we expect recoil after laser ablation to happen on faster timescales than vertex rearrangements. It also helps us to keep track of the length of the ablated edge, since it will not be involved in any vertex rearrangements. After each laser ablation, we evolve the tissue for another 150 time units, which is sufficiently long to allow the tissue to settle into a static equilibrium. Cells do not divide during this duration.

We analyse the utility of laser ablation summary statistics in inferring Λ and Γ in Figure 4. Specifically, we investigate the extent to which parameters can be inferred using the recoil distance, area asymmetry, and perimeter asymmetry. Each of these summary statistics is obtained from one laser ablation per parameter set. The latter two asymmetry measures were used by Farhadifar et al. [5] to constrain their parameter choices in manual parameter fitting. The area asymmetry measures how much the combined areas of the cells adjacent to the ablation change in comparison to changes in the length of the ablated edge. The perimeter asymmetry similarly considers changes in the combined cell perimeter of the two cells. Both of these measures quantify whether the cell shape changes upon laser ablation are limited to an extension of the ablated edge, or whether the cells respond as a whole by changing their total areas or perimeters. Similar to our previous observations of summary statistics of cell packing, the posterior distributions corresponding to summary statistics from laser ablations shown in Figure 4 have mean values far away from the reference parameter set, and a high degree of uncertainty is associated with these parameter estimates. For example, the posterior estimated using the ‘Area asymmetry’ summary statistic has a mean value of (Λ, Γ) = (-0.33, 0.11) and marginal standard deviations of 0.33 and 0.06 in Λ and Γ, respectively.

Note that Farhadifar *et al.* [5] restricted their parameter analysis to positive values of Λ, since the vertex distances always increased after the ablation, rather than contracted, indicating that the edges are under tension. Here, of the 88,479 parameter points for which we successfully conducted laser ablation experiments, only 982 lead to a contraction of vertex distances after cutting, illustrating that increasing vertex distances after ablation are common even when Λ is negative. Hence, we cannot confirm that positive vertex recoils dictate positive values of Λ.

In conclusion, we observe high parameter uncertainty for parameter estimates generated using each summary statistic in Figure 4. This suggests that the model parameters may be practically unidentifiable due to related roles of these parameters in the model, or that the amount of data used to calculate the summary statistics is insufficient to reliably infer model parameters.

### 3.2 Repetition of experiments reduces parameter uncertainty

In Figure 4 we used summary statistics from single snapshots of tissues containing approximately 500 cells. To test whether better parameter estimates can be achieved if more data are used we obtain posterior distributions by calculating summary statistics from multiple experiments. Specifically, we averaged all summary statistics of cell shapes introduced in Table 2 and Figure 4 across ten tissues generated using our model and the reference parameter set. Summary statistics from laser ablation experiments are averaged over 60 laser ablations, i.e. six laser ablations are conducted on each simulated tissue. To ensure that the laser ablations do not impinge on each other or on the tissue boundary, we pick one cell interface at the centre of the tissue in addition to five cell interfaces that are evenly distributed along a circle sharing the tissue centre and with a radius of a quarter of the tissue width.

The posterior distributions from all summary statistics displayed in Figure 5 have lower uncertainty and provide better parameter estimates than those in Figure 4. Specifically, the posterior distributions from the ‘Polygon number’ and ‘Area ratios’ summary statistics lead to parameter estimates close to the reference parameters, with (Λ, Γ) = (0.06, 0.05) and (Λ, Γ) = (0.03, 0.06), respectively. Note, that we use the reference parameter set Λ_obs_, Γ_obs_) = (0.12, 0.04) to generate observed data unless stated otherwise. The ‘Polygon number’ and ‘Area ratios’ summary statistics are vector-valued summary statistics of cell shapes. In contrast, one-dimensional first-order summary statistics of cell shapes do not lead to improved parameter estimates in Figure 5 over those in Figure 4. The corresponding posterior distributions have similarities with those obtained from single experiments. In particular, the posterior distributions generated using the one-dimensional summary statistics of cell shapes in Figure 4 appear again to sit along a line in parameter space. We conclude that vector-valued summary statistics that average properties of cells of different polygon classes separately perform better in estimating parameters than summary statistics averaged across all cells in the tissue.

**Figure 5:**
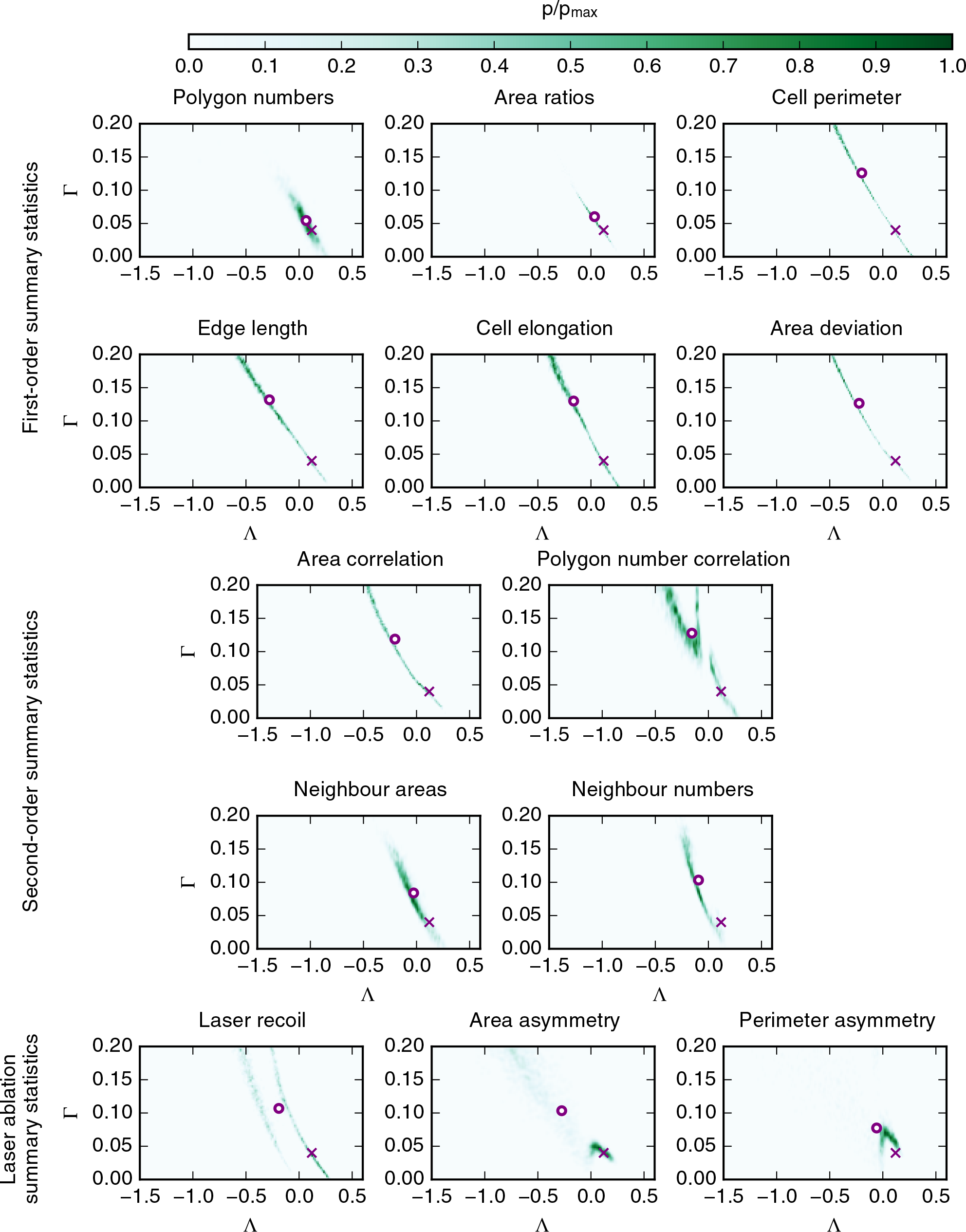
Posterior distributions of (Λ, Γ) obtained from different summary statistics of repeated experiments. Posterior distributions obtained using first-order summary statistics, second order summary statistics, and laser ablation experiments are shown in green. Parameter values are listed in Table 1 and summary statistics defined in Table 2. Crosses mark the reference parameter set used to generate the observed data. Circles mark the means of the posterior distributions. Posterior distributions were calculated using summary statistics averaged from ten experiments.

Second-order statistics of cell shapes obtained from multiple experiments appear to result in posterior distributions that are associated with high uncertainty for both one-dimensional or multi-dimensional summary statistics, and the associated parameter estimates lie far away in parameter space from the reference parameter set (Figure 5). This suggests that the data gathered across ten experiments at each parameter point are not sufficient to obtain reliable parameter estimates from second-order summary statistics of cell shapes. This reflects previous findings that second-order statistics tend to be associated with higher noise than first-order statistics [34].

Parameter estimates generated using summary statistics of multiple laser ablation experiments have lower uncertainty than those generated using summary statistics of single ablation experiments. However, the parameter estimates still lie far away from the reference parameter set, for example (Λ, Γ) = (-0.28, 0.1) in case of the ‘Area asymmetry’ summary statistic. The posterior distribution obtained from the ‘Laser recoil’ summary statistic is bimodel in the Λ direction, which may be an artefact of the boundary condition or of the normalisation of the recoil by the square root of the average tissue area. The posterior distributions from the ‘Area asymmetry’ and ‘Perimeter asymmetry’ boundary conditions contains mass in the vicinity of the reference parameter set, with maximum likelihood estimates at (Λ, Γ) = (0.09, 0.05) and (Λ, Γ) = (0.03, 0.07), respectively. This means that for these summary statistics the maximum likelihood provides a better parameter estimate than the mean of the distribution.

In summary, parameter estimates obtained by averaging summary statistics across multiple experiments in Figure 5 have lower uncertainty than those obtained from single experiments in Figure 5. However, a careful choice of summary statistic is important, since only few of the summary statistics in Figure 5 lead to reliable parameter estimates. Specifically, the vector-valued summary statistics ‘Area ratios’ and ‘Polygon numbers’ lead to parameter estimates close to the reference parameter set.

### 3.3 Posterior distributions are robust to the total number of samples and the number of accepted samples

When using ABC to infer posterior parameter distributions, it is important to check that the method is applied correctly. It is necessary to check that the total number of samples, *N*, is sufficiently large, and that the number of accepted regression samples, *M*, is sufficiently small in order to generate accurate representations of the posterior. In Figure 6 we compare the posterior distributions obtained with *N* = 100, 000 and *M* = 2, 000 (Figure 6A), to those obtained with *N* = 50, 000 and *M* = 1, 000 (Figure 6B), and those obtained with *N* = 100, 000 and *M* = 1, 000 (Figure 6C) for the summary statistics ‘Polygon numbers’ and ‘Area ratios’ in the case that multiple experiments have been conducted, as in Figure 5. In Figure 6B, both *N* and *M* are reduced while fixing the acceptance ratio *M/N* at 0.02. As Figure 6 shows, the posterior distributions obtained using the summary statistics ‘Polygon numbers’ and ‘Area ratios’ are not strongly affected by changes of the inference parameters *N* and *M* and the characteristics of these distributions, including their mean value, are preserved. For example, for the ‘Polygon numbers’ summary statistics, the mean values of (Λ, Γ) are (-0.066, 0.054), (0.080, 0.051) and (-0.060, 0.057) for cases A, B and C in Figure 6, respectively. We conclude that, for our choice of *N* and *M*, the ABC method has converged and yields accurate approximations of the posterior distributions.

**Figure 6:**
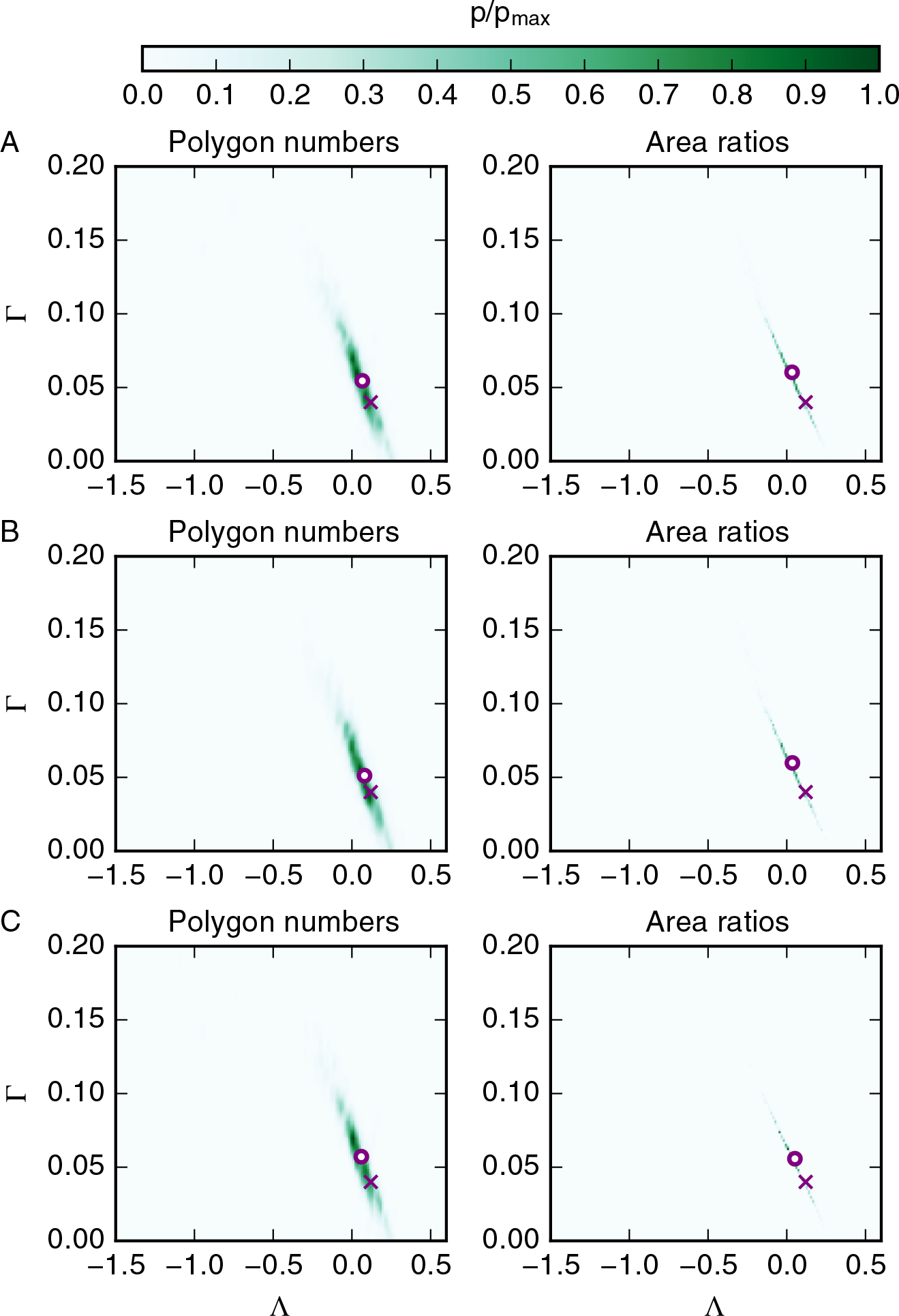
Posterior distributions are robust to inference parameters *N* and *M*. (A) Posterior distributions obtained using the summary statistics ‘Polygon numbers’ and ‘Area ratios’, the inference parameters *N* = 100, 000 and *M* = 2, 000 and ten experiments. (B) Posterior distributions obtained by halving the total number of samples, *N* = 50, 000. We also halve the acceptance threshold to *M* = 1, 000 in order to ensure that the acceptance ratio *M/N* is unaffected. (C) The posterior distributions obtained by halving the acceptance ratio, i.e. *N* = 100, 000 and *M* = 1, 000. Model and inference parameters are listed in Table 1 and mathematical descriptions of the summary statistics are provided in the text and listed in Table 2. Crosses mark the reference parameter set used to generate the observed data. Circles mark the means of the posterior distributions.

### 3.4 Selected combinations of summary statistics can further improve parameter estimates

After inferring the mechanical parameters of the vertex model using single summary statistics of cell packing and from laser ablation experiments, we investigate whether the parameter estimation process may be improved by combining summary statistics. We combine summary statistics by creating vector-valued summary statistics; if at least one of the summary statistics is already vector-valued, the vector is simply extended with entries for the other statistic. As described in Section 2.2, the individual entries of the combined, vector-valued summary statistics are rescaled by their respective standard deviations.

The posterior distributions resulting from three combinations of summary statistics are plotted in Figure 7. First, we combine the laser ablation summary statistics ‘Laser recoil’ and ‘Area asymmetry’ to test whether this combination leads to a posterior that is concentrated around the reference parameter values, since both the individual posterior distributions have high values in this region. However, the posterior distribution obtained using the combined summary statistic is instead widespread and leads to a parameter estimate far away from the reference parameter set, similar to the posteriors from either individual summary statistic. Next, we combine the ‘Area ratios’ and ‘Cell elongation’ summary statistics to test whether the contributions from the ‘Cell elongation’ summary statistic may help in ‘shifting’ the parameter estimate of the ‘Area ratios’ statistic closer to the reference parameter value. This is indeed the case, with (Λ, Γ) = (0.1, 0.05). Finally, we combine the ‘Area ratios’ summary statistic with the ‘Area asymmetry’. This is motivated by Farhadifar et al. [5], who used summary statistics of cell packing in combination with laser ablation experiments to estimate vertex model parameters. This combination of summary statistics does not lead to a more accurate parameter estimate (posterior mean) than the ‘Area ratios’ summary statistic, with (Λ, Γ) = (0.04, 0.06).

**Figure 7:**
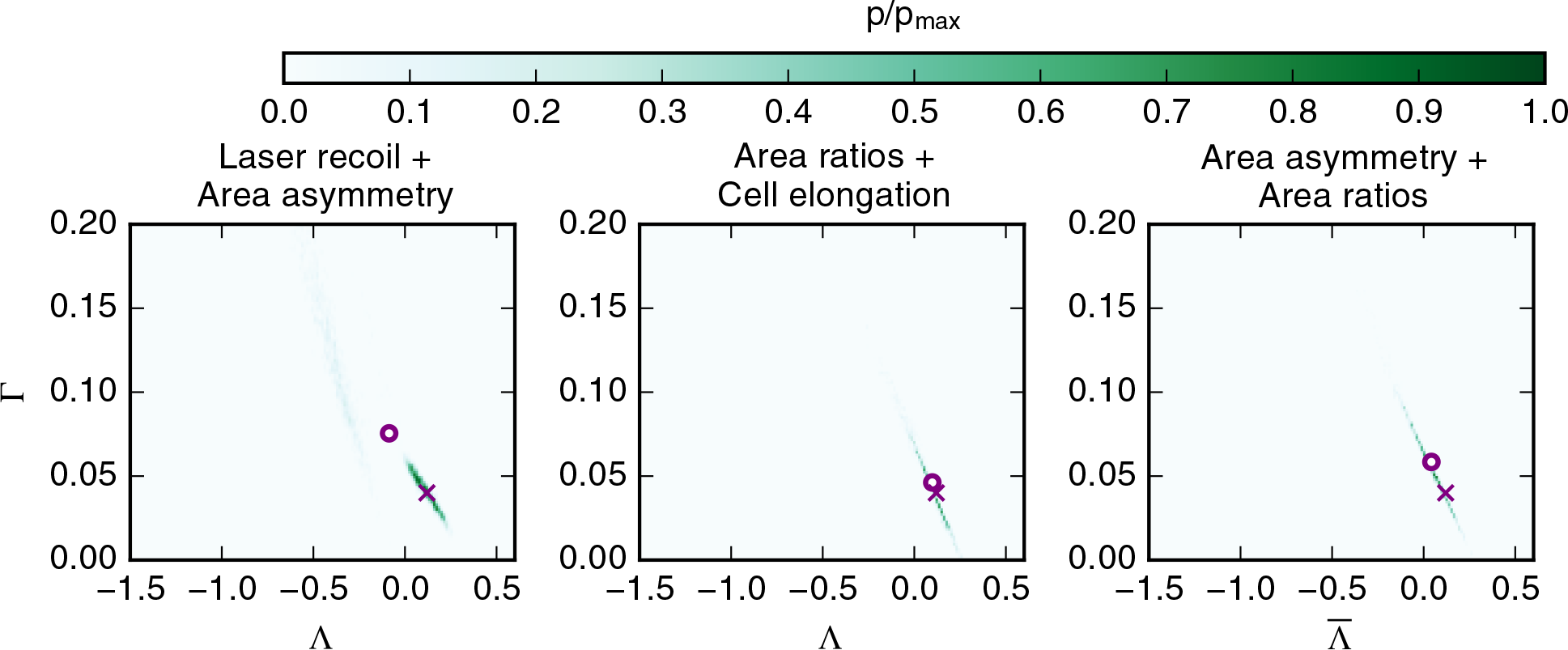
Combinations of summary statistics can lead to improved parameter estimates. Posterior distributions obtained when combining summary statistics are shown in green. The method for combining summary statistics is explained in the text. Model and inference parameters are listed in Table 1 and mathematical descriptions of the summary statistics are provided in the text and listed in Table 2. Crosses mark the reference parameter set of the data for which the posterior is estimated. Circles mark the means of the posterior distributions. Summary statistics are collected as averages from ten experiments.

Among all the summary statistics we considered, the most accurate parameter estimates are achieved when using the ‘Area ratios’ summary statistic in combination with the ‘cell elongation’ summary statistic; this combination performs slightly better than the ‘Area ratios summary statistic alone. A key benefit of using these two summary statistics is that they do not require laser ablation experiments to be conducted, i.e. they can be experimentally measured using non-invasive imaging methods.

### 3.5 Characteristics of posterior distributions are preserved for different reference parameters

We next test to what extent the ‘Area ratios’ summary statistic (Figure 8A) and its combination with the ‘Cell elongation’ summary statistic (Figure 8B) can be used to infer parameters of sample simulations with reference parameter sets elsewhere in parameter space, namely (Λ_obs_, Γ_obs_) = (-0.5, 0.1) and (Λ_obs_, Γ_obs_) = (-0.2, 0.15). For the reference parameter set (Λ_obs_, Γ_obs_) = (-0.5, 0.1), the parameter estimates (posterior mean) are further away from the reference value than for the previously analysed parameter point, with (Λ, Γ) = (-0.71, 0.14) in case of the combined summary statistic, and the posterior distributions are more widespread.

**Figure 8:**
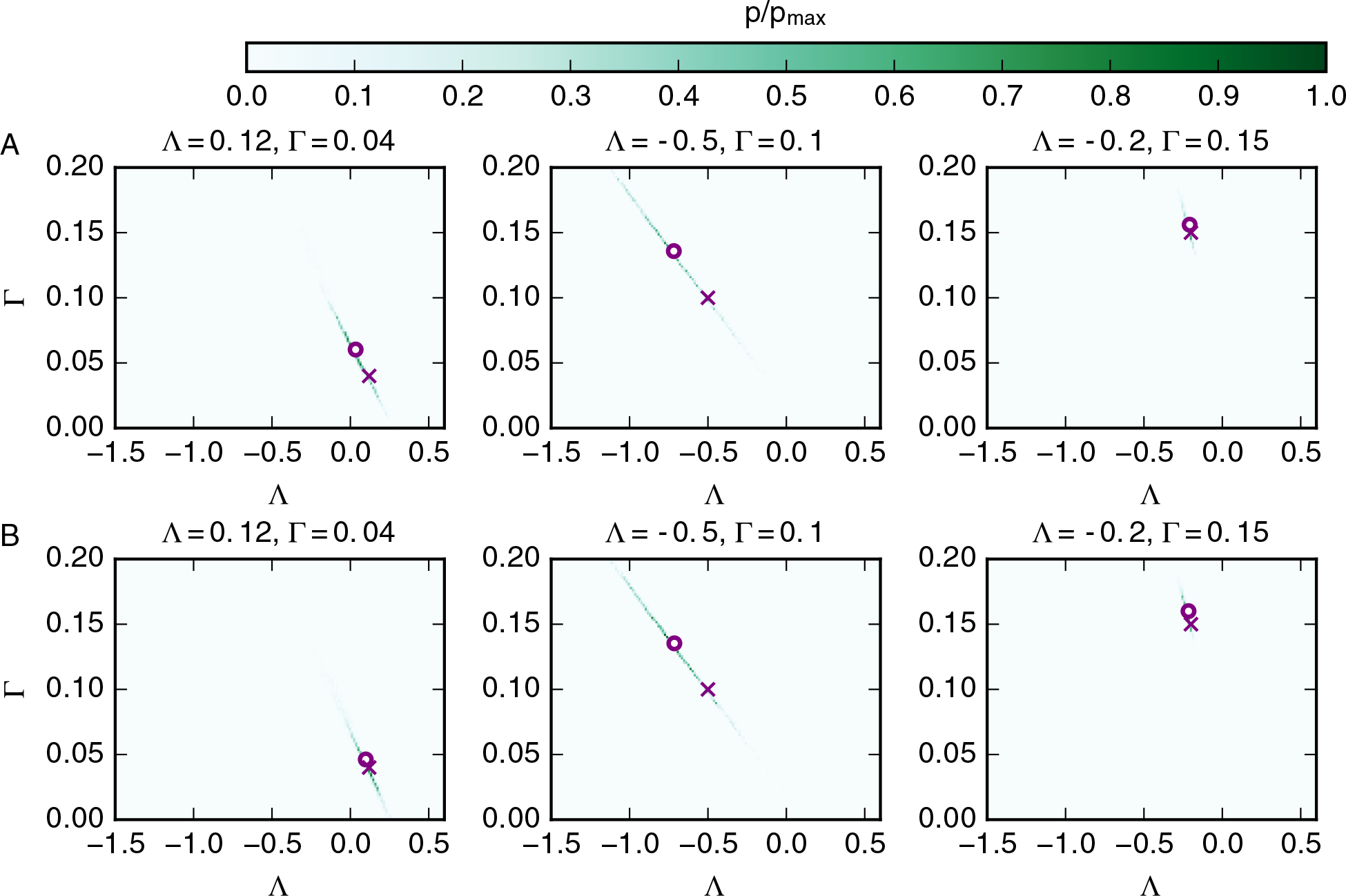
Characteristics of posterior distributions are preserved for different reference parameters. Posterior distributions obtained using the ‘Area ratios’ summary statistic (A) and the combination of the ‘Area ratios’ and ‘Cell elongation’ summary statistics (B) are shown in green. The method for combining summary statistics are explained in the text. Model and inference parameters are listed in Table 1 and mathematical descriptions of the summary statistics are provided in the text and listed in Table 2. Crosses mark the reference parameter of the data for which the posterior is estimated. Circles mark the means of the posterior distributions.

For the reference parameter set (Λ_obs_, Γ_obs_) = (-0.2, 0.15), the reference and the target parameter practically coincide with (Λ, Γ) = (-0.22, 0.16) and the parameter uncertainty is similar to the parameter uncertainty at (Λ_obs_, Γ_obs_) = (0.12, 0.04), with marginal standard deviations of 0.24 and 0.04 in Λ and Γ, respectively.

We conclude that posterior distributions obtained from *in silico* data generated using different reference parameter values share key properties. In particular, the accuracy and uncertainty of the parameter estimates are comparable.

### 3.6 Parameter estimates depend on the observed data at the reference parameter

In Figures 5, 7 and 8, accurate parameter estimates were achieved by a careful choice of summary statistic and ensuring that sufficient amounts of data were used. However, since the data generated by the model are inherently noisy, different parameter estimates may be obtained if different samples are used. In order to investigate how strongly the parameter estimate depends on the observed data we plot the parameter estimates obtained from ten different *in silico* data sets generated using the reference parameter set (Λ_obs_, Γ_obs_) = (0.12, 0.04) in Figure 9, using the ‘Polygon numbers’ summary statistic, the ‘Area ratios’ summary statistic and the combination of the ‘Area ratios’ summary statistic with ‘Cell elongation’. We note that each of the *in silico* data sets itself contains data from ten independent simulations.

**Figure 9:**
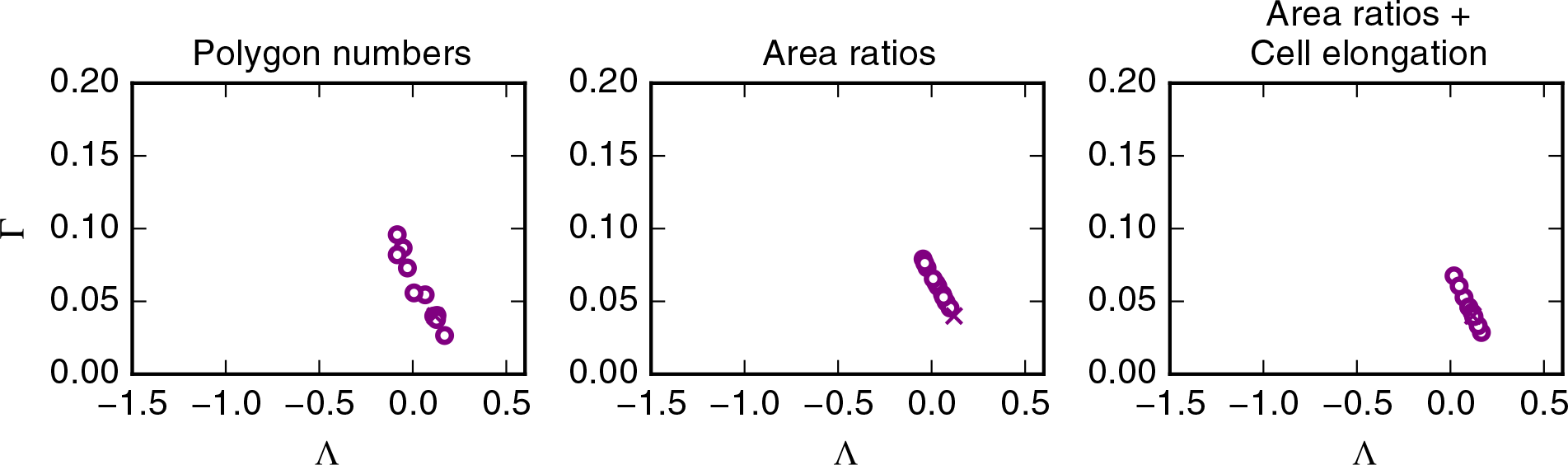
Parameter estimates depend on the observed data at the reference parameter. Circles mark the parameter posterior means obtained using the ‘Polygon numbers’ summary statistic, ‘Area ratios’ summary statistic, and the combination of the ‘Area ratios’ and ‘Cell elongation’ summary statistic evaluated for ten different realisations of the model at the reference parameter set (Λ_obs_, Γ_obs_) = (0.12, 0.04). The reference parameter set is marked by a cross. Model and inference parameters are listed in Table 1 and mathematical descriptions of the summary statistics are provided in the text and listed in Table 2.

The estimated parameter varies for the different reference data sets. The variation is wider for the ‘Polygon numbers’ summary statistic than for the ‘Area ratios’ summary statistic or the combined summary statistic, suggesting the ‘Polygon numbers’ summary statistic varies more between samples than the other summary statistics. All estimates (posterior means) using the ‘Area ratios’ summary statistic have larger values in Γ than the reference parameter set, which is not the case for the ‘Polygon number summary statistic’ or the combined summary statistic. Similar to the posterior distributions corresponding to these summary statistics, the estimated parameters lie on a line in parameter space, reflecting the fact that the parameters Γ and Λ regulate similar force contributions in the vertex model.

The data in Figure 9 confirm our observation that the combination of ‘Area ratios’ and ‘Cell elongation’ performs better than other summary statistics. In particular, the inferred parameter values vary less with the observed data than those inferred using the ‘Polygon number’ summary statistic, and inferred parameters lie on both sides of the reference parameter set, which is in contrast to parameter estimates from the ‘Area ratios’ summary statistic. For this combined summary statistic, the parameter estimates (posterior means) vary between (Λ, Γ) = (0.17, 0.03) and (Λ, Γ) = (0.02, 0.07).

## 4 Discussion

In this study, we have investigated the utility of a range of summary statistics in identifying the mechanical parameters of a vertex model of a proliferating epithelial tissue. For the first time, we provide uncertainty estimates arising from the use of such summary statistics by using ABC. We find that model parameters can be inferred from static images of cell shapes if summary statistics are carefully chosen and if sufficient amounts of experimental data are acquired. In particular, we identify that reliable parameter estimates can be achieved if a summary statistic is used that combines the mean area for cells of each polygon class with the average elongation of all cells in the tissue. Vertex model parameters can be inferred if this summary statistic is averaged across ten tissues containing approximately 500 cells. If less data are used, for example if summary statistics are only calculated from one sample tissue, then the uncertainty is large and parameter estimates are not reliable.

We find that summary statistics from invasive methods, such as laser ablation experiments, do not lead to better parameter estimates than summary statistics of cell shapes from static images. Similarly, second-order cell-shape statistics that characterise relationships of adjacent cells do not lead to better parameter estimates than first-order statistics, possibly reflecting higher noise in the second-order statistics if the same amounts of data are used. We identified that posterior distributions are robust to changes in the total number of samples and in the acceptance ratio, and that characteristics of the posterior distribution are preserved if different reference parameters are used. Further, we quantified the extent to which the parameter estimates depend on the observed data generated using the reference parameter set.

Summary statistics of cell shape have previously been used to constrain vertex model parameter space by Farhadifar *et al.* [5], who found that large regions of parameter space could give rise to experimentally observed values for such summary statistics. It has since been a common procedure to only use, for example, the distribution of cell neighbour numbers to select parameter values. Other summary statistics of cell shape have since been proposed, for example the cell elongation (or circularity) [12]. Here, we confirm previous findings by Farhadifar *et al.* [5] that high uncertainty may be associated with classical summary statistics of cell packing if insufficient data are used for parameter estimation. If sufficient data are used then summary statistics of cell packing may lead to reliable parameter estimates, in particular if vector-valued summary statistics are used that average cell properties over cells of each polygon class separately.

We further analysed, for the first time, whether second-order statistics of cell shapes, such as the correlation between areas of adjacent cells, can lead to improved parameter estimates. However, these second-order statistics do not lead to more accurate parameter estimates than summary statistics from first-order statistics of cell packing. Both types of summary statistics may suffer a similar drawback, namely a spread of the posterior distribution along an approximately linear line in parameter space, and similar posterior distributions were observed for summary statistics from laser ablation experiments. Summary statistics from laser ablation experiments lead to better parameter estimates if the maximum likelihood is used as parameter estimate instead of the average of the distribution.

The strong correlation between the parameters Λ and Γ in the posterior distributions corresponding to some summary statistics, such as the mean length of cell-cell interfaces or the standard deviation of cell areas, suggests that the model parameter space may be reduced to a lower-dimensional parameter space or transformed to a space of independent parameters. Previous studies have proposed reformulating equation (4) in terms of Γ and *p*_0_ = -Λ/2Γ [29, 35]. Estimated posterior distributions of (Γ, *p*_0_) are plotted in supplementary Figure S2, showing that parameter estimates of (Γ, *p*_0_) are associated with high uncertainty, similar to parameter estimates of (Γ, Λ). This high parameter parameter uncertainty in (Γ, *p*_0_) is to be expected, since this parameter transform does not reduce the parameter space. The transformed energy equation still depends on both parameters, Γ and *p*_0_, and since Γ and *p*_0_ are by definition interdependent, this transformation does not introduce uncorrelated parameters. Future work is required to understand to what extent the energy equation (4) may be reparameterised to expedite parameter estimation.

Parameter estimates from the vertex model using combinations of first order summary statistics are close to the ‘true’ reference parameter set. We see a decrease in parameter uncertainty when using ten experiments for parameter inference instead of one. Instead of simulating multiple tissues, it may also be possible to gain parameter estimates from cell packings of single, larger tissues. Here, we chose to simulate multiple small tissues since this facilitates parallelisation by ensuring that individual simulation runs are short. In future efforts, it may be possible to further reduce the parameter uncertainty by conducting more ‘experiments’ per parameter point. We did not investigate a further increase in the number of experiments per parameter point beyond ten due to prohibitive computational costs. Since posterior distributions obtained from summary statistics of laser ablation experiments have maxima close to the reference parameter values, future efforts may achieve better parameter estimates if summary statistics from more laser ablation experiments are used. Here, we used 60 laser ablations to calculate summary statistics and did not increase this number further due to prohibitive computational costs. For comparison, Farhadifar et al. [5] conducted 24 laser ablation experiments *ex vivo*.

When simulating laser ablation experiments, we set the line tension of the ablated edge, as well as the perimeter contractility of the adjacent cells, to zero. This procedure has originally been proposed by Farhadifar et al. [5]. Canela-Xandri et al. [10] also set the line tension of the ablated edge to zero, but remove the perimeter contractility contribution only from the ablated edge and not the remaining edges of the adjacent cells. More research is required to determine which method to implement laser ablations *in silico* is the most biologically realistic. This choice of implementation may have an effect on the resulting parameter estimates.

Laser ablations are a common procedure to measure the mechanical properties of a tissue *in vivo*. Other ways of tissue manipulation exist, for example through stretching or compressing the tissue [13]. Future work may include applying ABC to this type of experiment and measure the uncertainty of the associated parameter estimation. Experimentally, a possible avenue for future work might be to develop corresponding methods of tissue manipulation that are applicable *in vivo*, for example through locally perturbed tissue growth.

Here, we analysed the packing geometry and response to perturbations of tissues that emerge after periods of uniform growth, such as in the *Drosophila* wing imaginal disc. However, packing geometries emerge from from a wide range of epithelia under different conditions. Future work may investigate whether other ways of generating tissue packing lead to similar packing geometries, and indeed if the same summary statistics, or even the same simulation procedure presented here, may be used to measure vertex model parameters in different tissues.

In the present work, we use rejection-based ABC and identified the mean area per polygon class in combination with the average cell elongation as a suitable summary statistic to constrain vertex model parameters. Using this summary statistic it may be possible to apply adaptations of ABC in order to implement more efficient methods that have lower sample rejection rates, for example Markov Chain Monte Carlo [27], sequential Monte Carlo techniques [28], or machine learning approaches [36]. Specifically, the simulation procedure used in this study is designed to minimise the time that is required to generate the tissue. Using this simulation procedure, the tissue does not evolve to its final configuration through slow, quasistatic growth. Instead, tissue growth occurs within a dynamic regime of the model. We confirmed that summary statistics of cell packing are similar to what has previously been reported. Using alternative inference methods, it may be possible to use quasi-static timescales in the sample simulations. When using a biologically realistic implementation of the growth phase of the tissue it may be possible to improve the parameter estimates obtained here by using dynamic data of tissue growth throughout the simulation, instead of focussing only at the final configuration [37].

We rescaled components of vector-valued summary statistics by their respective standard deviations. This is a common procedure when conducting parameter inference, since it minimises the impact of summary statistics of high variability on the parameter estimate [21, 22]. However, the optimal choice for weights when combining summary statistics for parameter estimation is a matter of active research [38–40]. Here, we considered a large number of summary statistics and combinations and identified the mean area per polygon class in combination with the mean cell elongation as a suitable summary statistic for parameter inference. It is unclear whether better parameter estimates could be achieved in our work if different weights were used in vector-valued summary statistics, or if different combinations were applied.

Future methods of investigating vertex model parameters may include measurements of the mechanical relaxation time 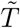. Measurement of the mechanical relaxation time has previously been proposed by [6] and [10]. This may, in turn, allow the precise implementation of cell cycle times in the vertex model.

Here, we have investigated parameter inference for vertex models propagated using the energy equation (4). Several other formulations have been proposed [3,41–43], including terms that take cell heights into account. To conduct parameter inference on vertex models using these altered energy equations, it would be necessary to conduct a preliminary analysis investigating the possible model behaviours, and the parameter regions where the models are physical. Future work may include using ABC for model selection on experimental data to investigate which of these energy equations provides the most plausible description of biological phenomena. Such work could, for example, include alterations to equation (4) that prevent mechanically-induced cell removal.

In our model implementation, vertex movement is deterministic and stochasticity occurs only through the random distribution of cell cycle durations. A more rigorous approach to include stochasticity in the model would be to include a noise term in the equation of motion (1), thus turning it into a Langevin equation [44]. Langevin equations are a common method to model particle motion on a microscopic scale. Future work is required to understand to what extent an additional noise term in the vertex movement equation may impact model behaviour or the utility of individual summary statistics in inferring model parameters.

The present work illustrates the importance of quantifying uncertainty when measuring parameters of stochastic models in biology. When few data are used for parameter inference, in our case if summary statistics are calculated from a single tissue containing 500 cells, it is difficult to identify a specific parameter set corresponding to a given tissue configuration. It follows that the exact choice of parameter values will not strongly influence the model behaviour and many modelling results will be robust to changes in the model parameters. Future efforts for parameter inference should include methods that go beyond measuring the non-dimensionalised parameters, and also encompass dimensional parameters such as the cellular stiffness 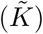, the target area of cells, the timescales of mechanical rearrangement, and parameters concerning cell cycle progression. Initial efforts in this direction have already been conducted [13, 15]. It will be important to extend such efforts to methods that quantify parameter uncertainty and are applicable in living tissues.

## Footnotes

1 Note that these boundaries are derived using free boundary conditions in the vertex model.

## Acknowledgements

The authors thank Ben Lambert for insightful discussions. JK acknowledges funding from the Engineering and Physical Sciences Research Council (grant number EP/N509711/1). REB is a Royal Society Wolfson Research Merit Award Holder and a Lever-hulme Research Fellow. AGF is supported by a Vice-Chancellor’s Fellowship from the University of Sheffield. The authors acknowledge the use of the University of Oxford Advanced Research Computing (ARC) facility in carrying out this work (http://dx.doi.org/10.5281/zenodo.22558).

